# A red algal polysaccharide influences the multicellular development of the choanoflagellate *Salpingoeca rosetta*

**DOI:** 10.1101/2024.05.14.594265

**Authors:** Olivia Perotti, Gabriel Viramontes Esparza, David S. Booth

## Abstract

We uncovered an interaction between a choanoflagellate and alga, in which porphyran, a polysaccharide produced by the red alga *Porphyra umbilicalis*, induces multicellular development in the choanoflagellate *Salpingoeca rosetta*. We first noticed this possible interaction when we tested the growth of *S. rosetta* in media that was steeped with *P. umbilicalis* as a nutritional source. Under those conditions, *S. rosetta* formed multicellular rosette colonies even in the absence of any bacterial species that can induce rosette development. In biochemical purifications, we identified porphyran, a extracellular polysaccharide produced by red algae, as the rosette inducing factor The response of *S. rosetta* to porphyran provides a biochemical insight for associations between choanoflagellates and algae that have been observed since the earliest descriptions of choanoflagellates. Moreover, this work provides complementary evidence to ecological and geochemical studies that show the profound impact algae have exerted on eukaryotes and their evolution, including a rise in algal productivity that coincided with the origin of animals, the closest living relatives of choanoflagellates.

## Introduction

The choanoflagellate, *Salpingoeca rosetta*, exemplifies the ability of unicellular eukaryotes to integrate complex environmental cues into their life history. Like many eukaryotes, nutrient availability drives life history transitions in *S. rosetta*^1^, including their sexual cycle^2^. Their main food source, bacteria, can produce another set of independent cues for choanoflagellates to detect, such as glycosyl lyases^,4^ that trigger mating and lipids^5,6^ that induce the development of multicellular rosettes through serial cell divisions (Fig. 1A). The roles of bacteria in the developmental transitions of choanoflagellates and their closest living relatives, the animals, have underscored that the evolution of both choanoflagellates and animals has been shaped by their intimate relationships with bacteria^7–10^.

**Figure 1:**
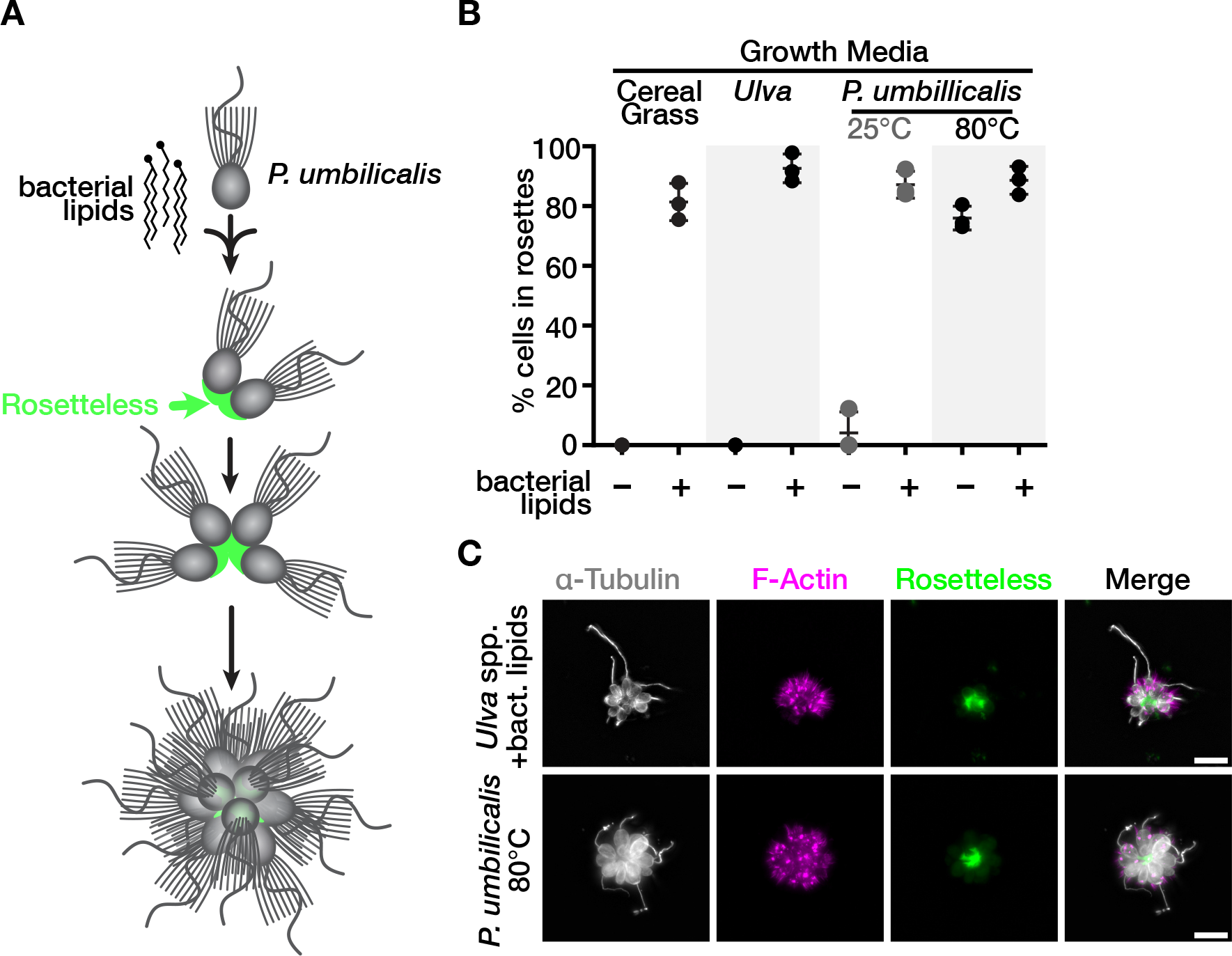
Growth media prepared from *Porphyra umbilicalis* induces multicellular development of the choanoflagellate *Salpingoeca rosetta* **(A)** *S. rosetta* senses environmental cues to develop into multicellular rosettes. Upon sensing lipids from the bacterium *Algoriphagus machipongonensis*, *S. rosetta* develops into rosettes through serial cell divisions while secreting an adhesive protein, Rosetteless (green), at the basal end of cells. Here we show that the red macroalga *P. umbilicalis* also induces rosette development. **(B)** A medium prepared from the red alga *P. umbilicalis* induces rosette development. We optimized preparation of media enriched with *P. umbilicalis* at low (25°C) and high (80°C) temperatures (Fig. S1). We compared rosette induction with (+) or without (–) bacterial lipids in media prepared from *P. umbilicalis,* the green alga *Ulva* spp., and Cereal Grass. Rosette induction was quantified by counting the number of single swimmers and cells in rosettes for a total of 500 cells. Independent triplicates of those counts (black dots) are shown with their mean and standard deviation (lines). Only media prepared from *P. umbilicalis* at 80°C induced rosette development in the absence of bacterial lipids delivered in outer membrane vesicles from *A. machipongonensis*. **(C)** *S. rosetta* secretes Rosetteless into the interior of multicellular rosettes induced with bacterial lipids or media enriched with *P. umbilicalis*. Immunofluorescent microscopy of rosettes induced with bacterial lipids or media enriched with *P. umbilicalis* visualizes the multicellular architecture of rosettes through an anti-alpha tubulin antibody (DM1A), which highlights, microtubules (grey), and phalloidin, which highlights filamentous actin (magenta). An antibody raised against Rosetteless (refs levin and rutaganira) shows the localization of this protein (green) in the interior of rosettes where the basal ends of cells meet. Scale bars denote 10 µm.

In marine environments, bacteria not only grow in the water column but also densely colonize surfaces of algae^11–15^. Macroalgae (also called seaweeds) harbor diverse microbial communities composed of both bacteria and microeukaryotes^16–18^. The interactions between these microbes and their macroalgal hosts is often specific. Some bacteria only grow on certain species of seaweeds, determined by enzymes that bacteria possess to break down polysaccharides that form algal cell walls^19–22^. For example, one bacterium, *Zobellia galactonorivans* colonizes *Porphyra umbilicalis*^23^, a red macroalga, and secretes an enzyme to degrade the excreted polysaccharide porphyran^19,20^. Similarly, microeukaryotes associate with particular types of macroalgae^16^ and take advantage of nutrients diffusing from macroalgae^24^.

Algae may have long served as attractors for diverse microbial communities, for geochemical evidence indicates that algae became more abundant during important epochs in eukaryotic evolution^25^, including the origin of animals^26^.

Fascinated by the influence that algae can exert on microbial communities and historical descriptions of choanoflagellates attached to algae^27–29^, we decided to test if we could prepare media from macroalgae to foster the growth of *S. rosetta* in the laboratory. In these experiments, we were surprised to find that *S. rosetta* developed into multicellular rosettes in media prepared from *P. umbilicalis* even in the absence of bacteria that produce rosette-inducing cues. Through a biochemical approach, we found that the inducing factor was porphyran, showing that polysaccharides from algae can serve as environmental cues for choanoflagellate life history transitions.

## Results

### *S. rosetta* develops into multicellular rosettes in media prepared from red algae

We initially set out to improve media for culturing *S. rosetta* by enriching seawater with macroalgae, for we previously observed that small alterations in media formulations increased the maximal cell density of *S. rosetta* in culture^30^. We reasoned that macroalgae may provide better nutritional support for the growth of *S. rosetta* and/or their bacterial prey, which is their obligate food source, because macroalgae accumulate high concentrations of essential nutrients from their environment^31–33^. Nutrients derived from macroalgae may also be more ecologically relevant for choanoflagellates because they and the bacteria that influence the life histories of choanoflagellates have previously been isolated from macroalgae^28,19,5^. Therefore, we prepared medias enriched with a red alga (*Porphyra umbilicalis*), a green algae (*Ulva* spp.), or a brown alga (*Saccharina latissima*) by steeping dried fronds of each macroalga in synthetic seawater (Table S1). After acclimating cultures of *S. rosetta* feeding on *Echnicola pacifica* to each media over three or more passages, we compared the proliferation of *S. rosetta* in media prepared from macroalgae to a traditional growth medium prepared from cereal grass^34,35^ (Fig. S1A). Medias prepared with *P. umbilicalis* and *Ulva* spp. both supported choanoflagellate growth as well as cereal grass media; whereas, *S. rosetta* grew more slowly and to a lower density in media prepared from *S. latissima*.

To our great surprise, we also observed that *S. rosetta* sporadically developed into multicellular, rosette colonies. Previous work found that rosettes only develop in the presence of inducing cues produced by bacteria, of which the most intensively studied are sulfonolipids from *Algoriphagus machipongonensis*, but the only bacterium in our culture was *E. pacifica,* which does not produce rosette inducing factors^2,6,36^. Intrigued by the infrequent appearance of rosettes in our cultures, we decided to optimize the preparation of red algae media for more reliable rosette induction (Fig. S1 and Table S1) and better cell proliferation (Fig. S2 and Table S2). In this optimization, media prepared at higher temperatures (Fig. 1B and Fig. S1B) or with a larger mass of *P. umbilicalis* (Fig. S1C) led to a higher proportion of *S. rosetta* developing into rosettes, but the same conditions did not result in rosettes developing in media prepared from *Ulva* spp. The absence of rosettes in media prepared from *Ulva* spp. or in media steeped with *P. umbilicals* at low temperatures is most likely due to the lack of inductive cues rather than deficient growth conditions, as the addition of rosette inducing factors from outer membrane vesicles of *A. machipongonensis* still caused rosette development in those medias.

We further confirmed that *S. rosetta* developed into rosettes by fluorescently staining colonies with an antibody that recognizes a secreted protein, Rosetteless, necessary for rosette development^36,37^. This antibody staining showed that rosettes grown in media prepared from *P. umbilicalis* contain Rosetteless at the basal ends of individual cells that point toward the interior of the rosette, just as in rosettes induced with bacterial cues (Fig. 1C). Altogether, these data show that media prepared from *P. umbilicalis* induces rosette development similarly to bacterial signals.

### A polysaccharide, porphyran, purified from *P. umbilicalis* induces rosette development

What factor in the media prepared from *P. umbilicalis* induces *S. rosetta* to develop into rosettes? We first thought the factor may come from a bacterium, for only bacterial cues have previously been described to induce rosette development^38,5,6,4,39^. Furthermore, *P. umbilicalis* can harbor the epiphytic bacteria *Z. galactanivorans* that induces rosette development^5^. To test if bacteria may have been the source of the inductive cue in our media preparation, we isolated bacteria from the dried fronds of *P. umbilicalis* that were used to make media. We found species from the genera *Microbacterium*, *Pseudomonas*, *Aureimonas*, and *Aeromicrobium* associated with the dried fronds. However, none of the isolated bacteria induced rosettes when added to cultures of *S. rosetta* feeding on *E. pacifica*. Although this negative result cannot completely exclude bacteria as the source of the inducing factor, we began to consider that the factor may come from *P. umbilicalis*. In support of this hypothesis, we also found that media prepared from other species of red algae, *Palmaria palmata* and *Chondrus crispus*, induced rosette development (Fig. S1D), indicating that the inducing factor may be a molecule commonly shared among those algae.

The observation that high temperature preparations of media more frequently induced rosettes encouraged us to biochemically purify a rosette-inducing factor from *P. umbilicalis* (Fig. 2A). Inspired by previous studies that identified rosette-inducing factors from bacteria^5,6^, we tested rosette induction for each step of the biochemical purification (Fig. 2B). After obtaining a soluble extract from *P. umbilicalis* with potent rosette induction, we first used a hydrophobic resin (silica functionalized with octadecyl moieties) to test if the inducing factor from *P. umbilicalis* was a lipid like the previously described bacterial cues, yet the rosette-inducing activity remained in solution after passing through the hydrophobic resin, indicating that the inducing factor from *P. umbilicalis* was hydrophilic and probably not a lipid. Next, we added a battery of enzymes to digest any proteins or nucleic acids that may have persisted in the purification and to degrade agar, which was making the extract more viscous. None of the enzymes, however, reduced rosette induction. We next wanted to know if the inducing factor was a large or small molecule, so we next dialyzed the extract with membranes that had a 40 kD molecular weight cutoff. The retention of rosette inducing activity in the dialysis bag indicated that the inducing factor was a macromolecule larger than 40 kD. In later purifications, we also varied the order of purification steps to place the hydrophobic resin at the end, which still produced a simplified extract that induced rosette development.

**Figure 2:**
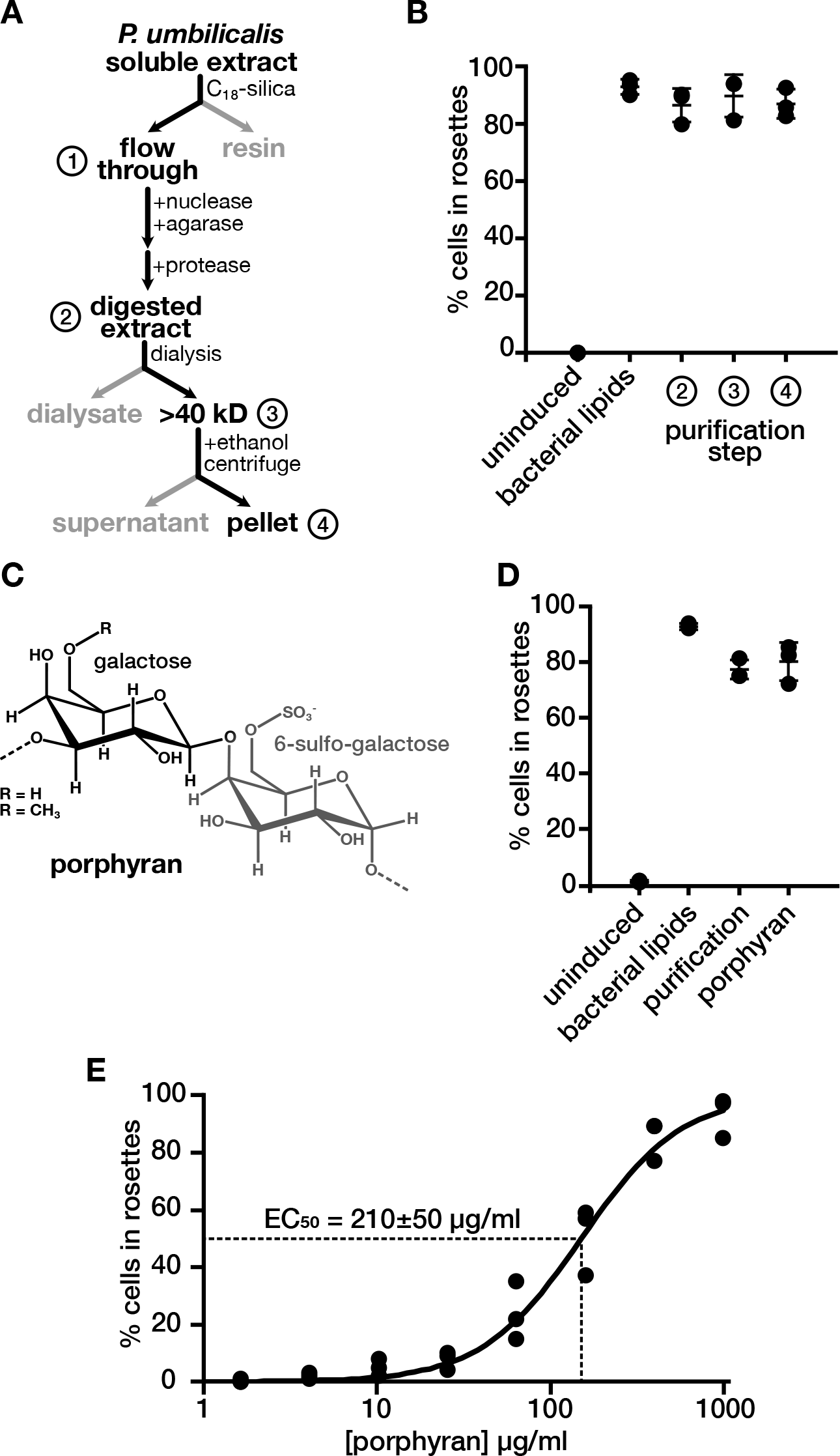
Porphyran, a sulfated galactan from *P. umbilicalis*, induces rosette development. **(A and B)** The purification of rosette inducing activity from *P. umbilicalis* extracts. (**A**) The purification scheme for rosette inducing activity started with the preparation of a soluble extract from *P. umbilicalis.* A hydrophobic resin (C_18_-silica) retained hydrophobic molecules while hydrophilic molecules passed through the resin in the ‘flow through,’ to which we added enzymes that degrade nucleic acids, agar, and proteins to produce the ‘digested extract.’ In this sample, we measured rosette inducing activity (as described in Fig. 1B) to confirm that the ‘digest extract’ retained rosette inducing activity **(B)**. Likewise, rosette inducing activity remained after dialyzing the ‘digested extract’ in a membrane with a 40 kD molecular weight cutoff. To this dialyzed sample, the addition of ethanol formed precipitates that were pelleted by centrifugation. The dissolved pellet retained rosette inducing activity. **(C)** The chemical structure of porphyran. Porphyran consists of linked galactose molecules with either a sulfonate or methyl functional group on the sixth carbon of alternating monosaccharides. **(D)** Porphyran induces multicellular rosette development. Porphyran purchased from a commercial source induces rosettes to the same extent as the purified rosette inducing activity from *P. umbilicalis* (labeled purification) and bacterial lipids from *A. machipongonensis*. Rosette induction was quantified as described in Figure 1B. **(E)** Rosette induction is directly proportional to the concentration of porphyran. Across a serial dilution, the effective concentration of porphyran that induces 50% of cells to form rosettes (EC50) is 210±50 µg/ml, an average calculated from fitting the hill equation (line) to independent triplicates of the dilution series (dots).

We reasoned that polysaccharides were the most likely type of macromolecule to still induce rosette development after the addition of enzymes and the depletion of hydrophilic molecule(s). After precipitating polysaccharides from the purified extract by adding ethanol, we found that molecules dissolved from the pellet induced rosettes (Fig. 2B). A carbohydrate analysis of this pellet showed that it was composed of galactose and sulfur (Fig. S3). This composition mirrors that of porphyran, a major extracellular polysaccharide from *P. umbilicalis* that is made of alternating subunits of galactose and 6-sulfo-galactose (Fig. 2C). To independently confirm that porphyran was the inducing molecule, we obtained porphyran from a commercial source and tested its ability to induce rosettes. This commercial source of porphyran induced rosettes to the same extent as our own preparation (Fig. 2D). We also characterized the potency of porphyran to induce rosette development in a dilution series, finding that the effective concentration for fifty-percent induction (EC_50_) was 210±50 µg/ml.

Assuming a minimum molecular weight of 40 kD for porphyran, this potency would correspond to a micromolar concentration of porphyran, which approaches the potency of rosette inducing lipids from *A. machipongonensis*^6^.

Because bacteria are an obligate food source for *S. rosetta*, we cannot yet discern if porphyran directly induces rosette development or acts through a bacterial intermediate. More mechanistic work is necessary to distinguish among these possibilities, yet we suspect that *S. rosetta* directly recognizes porphyran. In support of this hypothesis, we purified outer membrane vesicles from *E. pacifica* that had been grown in rosette inducing media, and those vesicles failed to induce rosettes when added to cultures of *S. rosetta* grown in non-inducing media (Fig. S4B). Moreover, the *E* pacifica genome^40^ lacks the family of glycosyl hydrolases for ingesting porphyran, and the addition of the constituent monosaccharides of porphyran to cultures of *S. rosetta* and *E. pacifica* failed to induce rosettes (Fig. S4A). Regardless of the exact induction mechanism, porphyran now provides a biochemical probe to further investigate how *S. rosetta* perceives this cue to induce rosette development.

## Discussion

The realization that red algae can influence the multicellular development of *S. rosetta* began with a search for improved media to grow choanoflagellates. Bacterivorous protists, like choanoflagellates, proliferate in enriched media that promote the growth of prey bacteria and deliver nutrients that neither protists nor their bacterial prey can produce^34^. For example, media prepared with lemon peel provides the limiting nutrient ubiquinone for the growth of the dinoflagellate *Oxyrrhis marina*^41^. We could only find one previous example of using macroalgae as a component in a growth medium^34^, so our work adds macroalgae to the repertoire of enrichments for protist growth media. To demonstrate the generality of macroalgae as media enrichments, we grew a variety of choanoflagellate species in media prepared from red or green algae (Table S3).

We identified porphyran as the molecule from red algal media that induces multicellular development in *S. rosetta.* This discovery provides a biochemical insight into the associations between choanoflagellates and algae. These interactions were evident in the earliest descriptions of choanoflagellates, in which algae were sometimes used as bait to collect choanoflagellates that can directly attach to macroalgae^27,28^. In those direct connections, we can now appreciate that algae may not only have been convenient landing pads for choanoflagellates but also a source of signaling cues that direct life history transitions. Proximity is likely a key factor for a freely swimming choanoflagellate to detect cues from algae and their epiphytic bacterial communities, for large macromolecules like proteins, vesicles, and porphyran only slowly diffuse from their initial source. Alternatively, large molecules could accumulate in enclosed environments such as ephemeral pools near coastlines. In fact, on a field trip near Waquoit Bay, Massachusetts, we found an example of a puddle on a jetty in which a piece of red algae laid (Fig. S5B). The water in this puddle was teaming (∼10^4^ cells/ml) with a choanoflagellate species that formed chain colonies (Fig. S5C). Similarly, splash pools in Curaçao support the growth of another colony-forming choanoflagellate, *Choanoeca flexa*^42,43^.

Algae may emit a variety of cues into their environment to mold their surrounding microbial community. For example, *Asterionellopsis glacialis*, a diatom, by altering the secretion of metabolites^44^. Small metabolites can also attract phagotrophic protists to graze on phytoplankton^45^. During phytoplankton blooms, diatoms secrete copious quantities of the polysaccharides laminarin^46^ and fucoidan^47^, which may provide food and/or environmental cues for bacteria and microeukaryotes. The liberation of porphyran and other polysaccharides from macroalgae, however, may often depend on epiphytic bacteria that can break down polysaccharides that form algal cell walls^20^, and seasonal^48^ and life history^49^ variations in polysaccharide modifications may make the degradation of cell walls more or less efficient. Tidal forces and desiccation between tides may further contribute to the release of those polysaccharides. All of these possible variations emphasize how a multiplicity of environmental factors can converge to alter the interactions within the habitats formed by algae. The work presented here emphasizes that *S. rosetta* is a suitable model system to investigate how the confluence of bacterial, algal, and physical factors influence the responses of microeukaryotes to complex environmental cues.

## Acknowledgements

We thank the following individuals for insights and support that helped advance this work: Susan Brawley (U Maine), Hilary Morrison (Marine Biological Laboratory), Nicole King (HHMI/UC Berkeley) and Flora Rutaganira (Stanford U) for generously sharing antibodies and choanoflagellate strains. We are grateful to Fredrick Leon, Vicki Deng, and Ben Larson for providing critical advice and feedback as well as supporting early stages of the project. We thank the Complex Carbohydrate Research Center at the University of Georgia for performing glycosyl composition analysis, which was supported by NIH Grant R24GM137782. This work was supported by awards to DSB from the Chan Zuckerberg Biohub, the UCSF Sandler Program for Breakthrough Biomedical Research, a David and Lucile Packard Foundation Fellowship in Science and Engineering, and a Marine Biological Laboratory Whitman Center Fellowship sponsored by Nikon Inc. GV was a fellow in the UCSF IMSD Fellow and supported by a training grant for the UCSF Tetrad Graduate Program Training Grant T32GM139786.

## Author contributions

OP, GVE, and DSB designed experiments, conducted research, and wrote the manuscript. DSB conceived of the project and obtained funding.

## Materials and Methods

### Preparation of seawater media enriched with algae (Table S1)

(Note: All recipes for preparing media are in Table S1)

Media for culturing choanoflagellates is prepared from a concentrated stock of artificial seawater enriched with algae. To prepare this concentrated stock, 10 g of dried algae fragments (Maine Coast Sea Vegetables) is homogenized in 1 l of artificial seawater (ASTM D 1141, Ricca Cat No. 8363-5) by stirring at 400-500 rpm on a plate stirrer for 3 h and at the following temperatures: 80°C to prepare rosette inducing media from *P. umbilicalis* or 40°C for general cell culture media with minimal rosette induction activity. Afterwards, we removed large debris by first filtering algae/seawater mixture through two layers of Miracloth (EMD Millipore, Cat No. 475855-1R) and then once through a standard coffee filter and then twice through a Buchner funnel lined Whatman Grade 1 Filter Paper (Cytivia, Cat No. 1001-090). Note that the coffee filter is optional but highly recommended to increase the speed of filtration through the buchner funnel. Finally, the crudely filtered extract is vacuum filtered through a sterile 0.22 µm polyethersulfone (PES) membrane connected to a sterile flask (Fisherbrand, Cat No. FB12566504). This final filtrate is called 100% Algae Stock.

We dilute the concentrated stock of algae into artificial seawater along with mineral, trace metal, and vitamin supplements to prepare a final media for cell culture. To do so we dilute stocks of 100% Algae Stock, 1000x (Potassium Iodide, Sodium Nitrate, Sodium Phosphate), 1,000x L1 vitamins^54^, and 1,000x L1 trace metals^54^ in artificial seawater to yield a final 1x concentration of each component. A 1,000x Sodium Silicate stock can also be added for organisms, like loricate choanoflagellates and diatoms, that require silicate for growth. After all of the components are combined, the media is vacuum filtered through a 0.22 µm PES filter connected to a sterile flask. The resulting media is called 25% Algae Media.

### Maintaining Cultures of *Salpingoeca rosetta*

Strains of *S. rosetta* were co-cultured with *Echinicola pacifica* bacteria (SrEpac; American Type Culture Collection [ATCC], Manassas, VA, Cat. No. PRA-390)^2^ in media enriched with dried cereal grass or seaweeds as described above and Table S1. To maintain cultures of rapidly proliferating cells at 27°C, we passaged *S. rosetta* from cultures that reached ∼10^5^ cells/ml into fresh media at dilutions of ∼1/50 (daily) or ∼1/100 (every two days). The timing and dilution factor were adjusted according to growth parameter measurements for each media condition (Table S2). Cultures were maintained in vented, treated culture flasks and in media volumes that corresponded to 0.24 ml per cm^2^ of culture flask surface area. To characterize the phenotypic effect of bacteria isolated from *P. umbilicalis* on *S. rosetta*, added the isolated bacteria to SrEpac and passed these cultures three times before assessing the level of rosette induction (as described below).

### Preparation of rosette inducing factors (Figures 1, 2, S1, and S4)

Outer membrane vesicles from *A. machipongonensis* (ATCC BAA-2233) or *E.pacifica* were prepared from the supernatant of bacterial cultures as previously described with a few modifications. First, bacteria were grown in different sources media prepared from *P. umbilicalis* or *Ulva* spp. Second, after harvesting outer membrane vesicles from the filtered supernatant by centrifugation for 3 h at 150,000 x g and 4°C, the membrane pellet was transferred to a tared microcentrifuge tube for determining the crude mass of the outer membrane vesicles. A buffer of 50 mM HEPES-KOH, pH 7.4 was added to the outer membrane vesicles to achieve a final concentration of 10 mg/ml. After resting overnight at 4°C, the solution of outer membrane vesicles was filtered through a sterile 0.45 µm polyethersulfone syringe filter and stored at 4°C for later use. Solutions of porphyran either purified from *P. umbilicalis* (see below) or purchased from Biosynth (Cat. No.YP157502) were resuspended in 50 mM HEPES-KOH, pH 7.4 to a final concentration of 20 mg/ml and then sterile filtered in a 0.45 µm polyethersulfone syringe filter. In the same manner, we prepared solution of D-galactose-6-O-sulphate (Biosynth, Cat. No. MG00761) and D-galactose (Fisher Bioreagents, Cat. No. G1-100).

### Rosette-Induction Assays (Figures 1, 2, S1, and S4)

Inducing cues were added to samples of SrEpac that were seeded at a density of 10^4^ cells/ml for *S. rosetta* in each well of a 12-well culture plate with 1 ml of culture per well. After 22-36 h from induction, cells were immobilized by adding 10 M LiCl to 500 µl sample of SrEpac for a final concentration of 500mM, and chains of slow swimmers or cellular aggregates were disrupted by vortexing. To quantify percent rosette induction, we counted a total of 500 cells (Bright-Line hemacytometer, Hausser Scientific) and scored them as single cells or cells within rosettes. Rosettes were defined as groups of four or more cells that resisted agitation (vortexing) and were centrally organized around their basal ends^6^. Each induction experiment was performed as independent triplicates.

### Quantifying the growth dynamics of *S. rosetta* (Figures S1 and S2 and Table S2)

SrEpac was inoculated into fresh media at a density of 10^4^ cells/ml for *S. rosetta*. The freshly inoculated cultures were distributed in 0.5 ml aliquots into each well of a 24-well plate. At each time point over a time course, one of the wells was homogenized and then fixed with the addition of 20 µl of 37.5% formaldehyde. This sample was then loaded into the hemocytometer (Bright-Line hemacytometer, Hausser Scientific) to determine the cell density. An independent triplicate was taken for each time point. We calculated the doubling time (*D*), maximal density (*M*), and lag time (*T*) for each time course by fitting the data with a least absolute deviation algorithm to a modified version of a logistic growth equation that explicitly models the lag time with a heaviside step function:

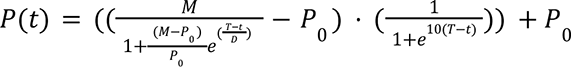

where *P(t)* is the cell density at a given time (t) and *P_0_* is the initial cell density.

### Immunofluorescent staining (Figure 1)

We adapted this method from one developed by Fredrick Leon^55^. A day before imaging SrEpac was seeded 10^4^ cells/mL of *S. rosetta* in media prepared from either *P. umbilicalis* or *Ulva* spp., which were supplemented with outer membrane vesicles from *A. machipongonensis*. The cells were grown at 27°C for one day. On the day of imaging, chambered glass coverslips (ibidi, Cat.No: 81507) were coated with 10mg/mL Poly D-lysine hydrobromide (MP Biomedicals, Cat. No. 102694) and then washed 3 times with 50uL of filtered artificial seawater (ASW). With a wide-bore pipette tip, 70 µl of rosettes were pipetted into a well of the coated coverslip and allowed to settle for 15 minutes. Afterwards, 50 µl of the liquid was removed, leaving 25 µl behind to avoid damaging delicate cell structures by drying the coverslip. The cells were first fixed by gently adding 50 µl of cytoskeleton buffer^56,57^ (10 mM MES, pH 6.1; 138 KCl, 3 mM MgCl_2_; 2 mM ethylene-glycol-bis(β-aminoethylether)-*N*,*N*,*N*’,*N*’-tetraacetic acid [EGTA]-KOH; 15% (w/v) sucrose) that also contained 3% paraformaldehyde (PFA). After incubating for 5 min at room temperature, 50 µl was removed. Cells were additionally fixed by adding 50 µl of the same fixation buffer that was now supplemented with a low concentration to Tween-20 (Cytoskeleton Buffer with 3% PFA and 0.07% Tween-20), which we found to improve the the preservation of cellular structures throughout subsequent step. Cells were incubated for 5 min at room temperature before removing 50 µl of the liquid. The fixation was quenched with the addition of 50 µl of quench buffer (Cytoskeleton Buffer with 0.3M glycine, pH 6.1). Immediately after, 50 µl was removed, and 50 µl of permeabilization buffer (LICOR Intercept PBS Blocking Buffer [LI-COR Biosciences Cat. No. 927-70001]; 8% (v/v) methanol, and 1% (v/v) Tween-20) was added to the coverslip and incubated for 30 minutes. Afterwards, wells were washed by replacing 50 µl of the liquid with antibody buffer (LICOR Intercept PBS Blocking Buffer with 1% (v/v) Tween-20), and we repeated this step for a total of two washes. Afterwards, 50 µl of an antibody mixture diluted in antibody buffer was added to the well. The antibodies were diluted as follow: 1:200 of 200 U/ml phalloidin-Alexa488, 1:500 mouse anti-αTubulin Monoclonal Antibody (DM1A) (Invitrogen, Cat. No. 62204), 1:1000 alpaca anti-mouse IgG1-Alexa555, 1:500 rabbit anti-Rosetteless (generously provided by Nicole King and Flora Rutaganira), and 1:1000 alpaca ant-rabbit IgG-Alexa647. The antibodies were incubated for 1 hour at room temperature and in the dark. Afterwards, the well was washed twice with 1x PEM (100 mM PIPES, 1 mM EGTA, 1 mM MgSO4, pH 6.9 with KOH) before imaging.

### Widefield Microscopy (Figure 1)

We visualized immunofluorescent samples on a Nikon Eclipse Ti2-E inverted microscope outfitted with a D-LEDI light source, Chroma 89401 Quad Filter Cube, and 60x CFI Plan Apo VC NA1.2 water immersion objective. Z-stacks of samples were imaged with two-fold binning (120 µm/pixel) on a Nikon Digital Sight 50 M Camera with 100-500 msec exposure times for each channel and variable illumination intensities to extend the dynamic range of signal without photobleaching samples. Images were processed in FIJI by projecting the maximal intensity of a central section of the rosette, subtracting the background (8 pixel, sliding paraboloid radius), and enhancing the contrast (saturating 0.2-0.4% of the pixels).

### Purification of rosette inducing activity from *P. umbilicalis* (Figure 2)

Dried algae was finely ground by flash freezing in liquid nitrogen and then grinding in a mortar and pestle chilled with liquid nitrogen. Into 200 ml of 50mM Tris-HCl, pH 8.0, 10 g of finely ground dried algae was stirred for 1 h at 400-500 rpm and 80°C. The homogenate was clarified by filtration through two layers of miracloth and then by centrifugation at 16,000 x g at 4°C for 40 minutes in a fixed angle centrifuge. Nucleic acids and agar were degraded in the clarified extracted by adding benzonase (35 units/ml, Thermo Fisher Scientific, Cat. No. 88700) and agarase (0.2 units/ml, Thermo Fisher Scientific, Cat. No. EO0461) and then incubating at 42°C for 1 hour. To degrade proteins, we treated the filtrate with pronase (10mg/mL, Sigma-Aldrich) for 1 hour at 37°C). Afterward, enzymes were heat-inactivated at 90°C for 20 minutes. The digested extract was dialyzed through a 20kD or 40 kD membrane (Slide-A-Lyzer, Cat. No. 66003) against ultrapure water for 24 h at 4°C . To deplete lipophilic contents, we flowed the dialyzed extract through a hydrophobic resin (Sep-pak Plus Short tC18, Waters Corp, Cat. No. 50-818- 646). To prepare the resin, we first passed a 2:1 ratio of methanol and chloroform, then equilibrated it in 50 mM Tris pH 8.0). To precipitate rosette inducing activity from the flow-through, 4 volumes of ethanol were added incubated overnight at -20°C. Afterward the pellet was recovered by centrifuging at 16,000 x g for 40 min at 4°C. The pellet was dried for 30 minutes at room temperature and then resuspended in 50mM Tris-HCl, pH 8.0 to 10 mg/ml. Samples were flash-frozen with nitrogen and stored at -80°C for later use.

### Carbohydrate Content Analysis (Figure nd S3)

Carbohydrate content of the purified pellet from *P. umbilicalis* extracts was performed by the Complex Carbohydrate Research Center Analytical Services at the University of Georgia.

The carbohydrate analysis was performed by derivatizing samples with *O*-trimethylsilyl (TMS) for characterization by gas chromatography/mass spectrometry (GC-MS)^58,59^. Lyophilized samples were hydrolyzed using 1 M methanolic HCl for 16 h at 80 °C and then cooled to room temperature before drying with N_2_ gas. 8 drops of methanol, 4 drops of pyridine, 4 drops of and acetic anhydride were added for N-acetylation, and the samples were heated at 100 °C for approximately 30 min. Once cooled, the samples were dried again, then 15 drops of Tri-Sil was added and incubated at 80 °C for 30 min. The samples were dried down one last time, before adding 150 μL of hexane to each. Each was briefly vortexed and centrifuged, then the contents of each sample transferred to GC-MS vials. 1 μL of each was injected into the GC-MS for analysis.

Sulfate analysis was performed by first creating a standard curve with a 1-mg/mL sodium sulfate (Na_2_SO_4_)/hydrochloric acid (HCl) solution (Fig. S3B). In addition, a 0.83-mg/mL solution of the sample was made with 200 μg (40 μL) of *Porphyra umbilicalis* from a 5-mg/mL sample solution, and 200 μL of 1.5 M HCl. The sample and standards were made in glass screw-cap tubes and incubated overnight at 80 ^°^C. A gelatin solution was made by dissolving 0.5 g gelatin and 100 mL of water at 60 ^°^C. Once dissolved, the solution was cooled to room temperature, and 2.5 mL of 6 M HCl and 0.5 g of barium chloride (BaCl_2_) were added and stirred/mixed with the gelatin solution overnight. The next day, the samples were removed from the heating block and cooled to room temperature. Once cool, 467 μL of the barium chloride-gelatin solution was added to 200 μL of each sample/standard. The solutions were vortexed to mix, and then 200 μL of each was added to a 96-well plate. The samples were allowed to incubate at room temperature for 10 minutes. The absorbance was read at 360 nm, blanking with the first point of the standard curve.

---

**Figure S1:**
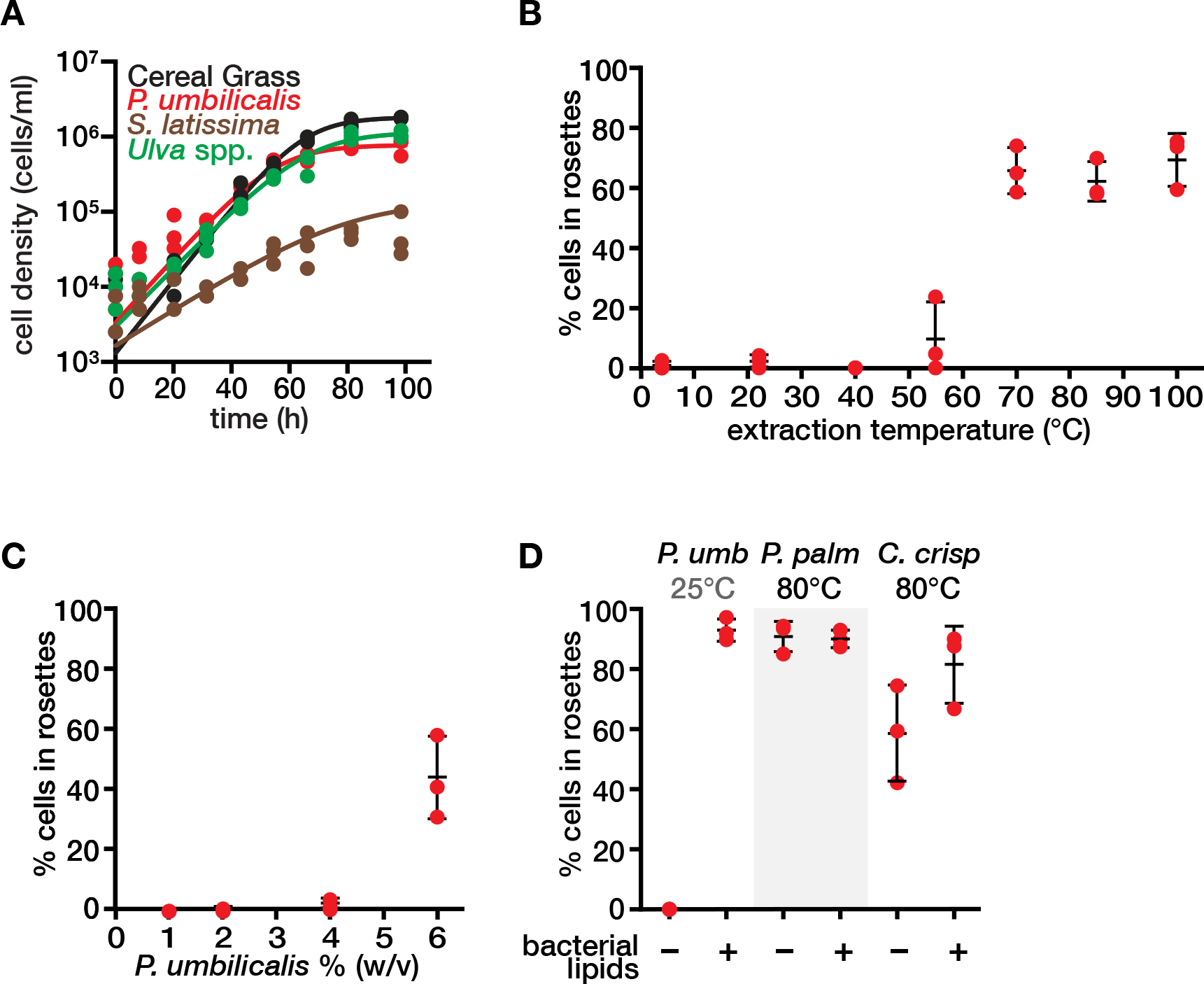
Growth media prepared from macroalgae support choanoflagellate growth and induce rosette development. **(A)** *S. rosetta* grows in media enriched with algae as well as media prepared with Cereal Grass. A media prepared from either *P. umbilicalis, Ulva Spp.,* or Cereal Grass media supports the growth of *S. rosetta,* and reaches its maximum carrying capacity of 10^6^ cells/mL at around 40 h post inoculation. In comparison, media from the brown alga *S. latissima* only minimally supported *S. rosetta* growth, reaching a significantly lower carrying capacity 10^5^ cells/mL) at 100 h post inoculation. **(B)** The rosette inducing activity from media prepared from *P. umbilicalis* increases with the temperature of extraction. Extraction of 1% (w/v) *P. umbilicalis* at temperatures between 10-50°C resulted in almost no rosette induction. In contrast, extraction temperatures above 55°C, resulted in ≥70% of S. rosetta cells developing into rosettes . Rosette induction was quantified as described in Figure 1B. **(C)** Preparing media with a greater mass of *P. umbilicalis* leads to increased rosette induction. To understand how the red algal infusion mass impacted rosette induction, we assessed rosette inducing activity of media prepared with 1, 2, 4, or 6% (w/v) *P. umbilicalis* at 25°C. Importantly, we diluted this media in proportion to the infusion mass to test rosette development under consistent nutrient levels. Without diluting the media in this manner, higher masses of *P. umbilicalis* in the media preparations resulted in a decrease in *S. rosetta* proliferation due to rapid *E. pacifica* growth. At this lower infusion temperature, only media made with 6% (w/v) of *P. umbilicalis* induced rosettes. Rosette inducing activity was quantified as in Figure 1B. **(D)** Media prepared from two additional species of red algae, *Palmaria palmata* and *Chondrus crispus*, induce rosette development. Seawater that was steeped at 80°C with 1% (w/v) of *P. palmata* or *C. crispus* yielded media that induced 60-80% *S. rosetta* cells to develop into multicellular rosettes. We compared this development to a negative control, in which seawater steeped at 25°C with 1% (w/v) of *P. umbilicalis* to produce a media that does not induce rosette development. Rosette induction was quantified as described in Figure 1B.

**Figure S2:**
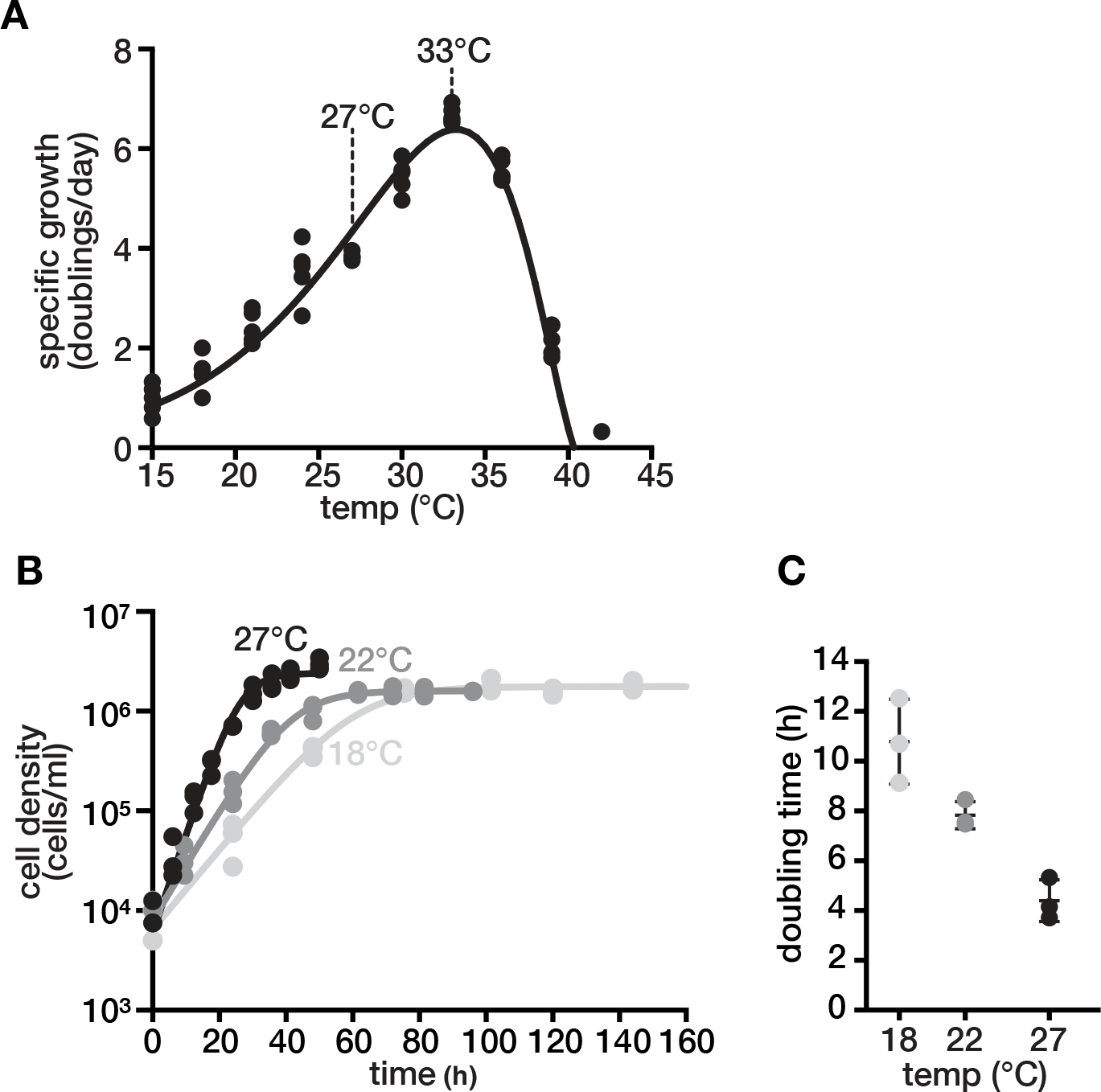
*S. rosetta* grows over a broad range of temperatures in media prepared from *P. umbilicalis*. **(A)** The specific growth of *S. rosetta* at varying temperatures. We seeded cultures with 10^4^ cells/ml of S. rosetta at the indicated temperatures and then counted the cell density after one day of growth. We converted cell density into specific growth (number of doublings per day) with the following equation: Specific Growth = log_2_(Cell Density•10^-4^). The data were fit to a model of temperature-dependent growth formulated by Blanchard^50,51^. From this curve, we numerically calculated first derivative to find the optimum temperature for specific growth at 33°C and the second derivative to find the inflection point in specific growth 27°C. The inflection point indicates the temperature at which the acceleration in specific growth as a function of temperature is zero and therefore robust to temperature deviations^52^. **(B and C)** *S. rosetta* grows rapidly at 27°C. In media prepared from *P. umbilicalis*, we acclimated *S. rosetta* to grow at 18°C, 22°C, or 27°C over three passages. Afterwards, we measured the growth parameters for *S. rosetta* over a time course (**B).** The maximal growth rate (also known as doubling time) was inversely proportional to temperature (**C**). Notably, *S. rosetta* growth almost twice as fast at 27°C in media prepared from *P. umbilicalis* than in standard media conditions at 22°C (Table S2).

**Figure S3:**
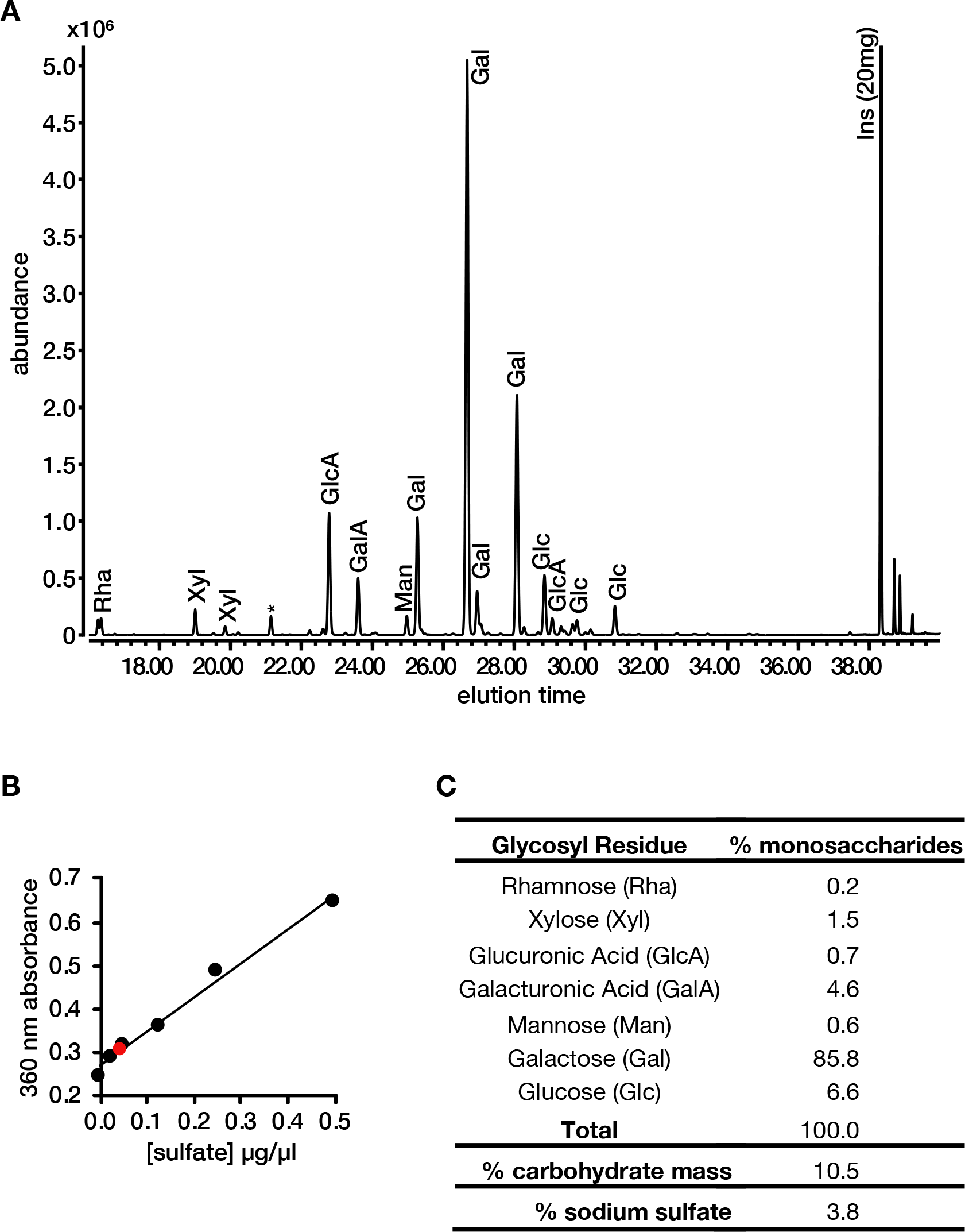
Galactose is the primary monosaccharide in carbohydrates purified from *P. umbilicalis*. (A) A chromatogram shows that galactose is the major monosaccharide in purified extracts from *P. umbilicalis*. The purification from *P. umbilicalis* was hydrolyzed with strong acid and derivatized for gas chromatography followed by mass spectrometry (GC-MS) to identify individual monosaccharides. The total area under the peaks that correspond to galactose accounts for ∼86% of the total monosaccharide abundance. (B) Sulfate quantification in the purified extract from *P. umbilicalis*. Sulfate concentration was determined with a gelatin-barium chloride method^53^. The method was calibrated with a standard curve (black dots and line) to determine the concentration of sulfate in our purified sample (red dot), showing that sulfate is a substantial component of the purified sample. (C) Table summarizing the abundance of components in the purified extracts. While a number of other sugar residues are present in the sample, galactose residues, likely from porphyran, make up 85.8% of the carbohydrate biomass. In addition, the product is 3.8% sodium sulfate, which points to the presence of the 6-sulfo-galactose residue of porphyran (Fig. 2C).

**Figure S4:**
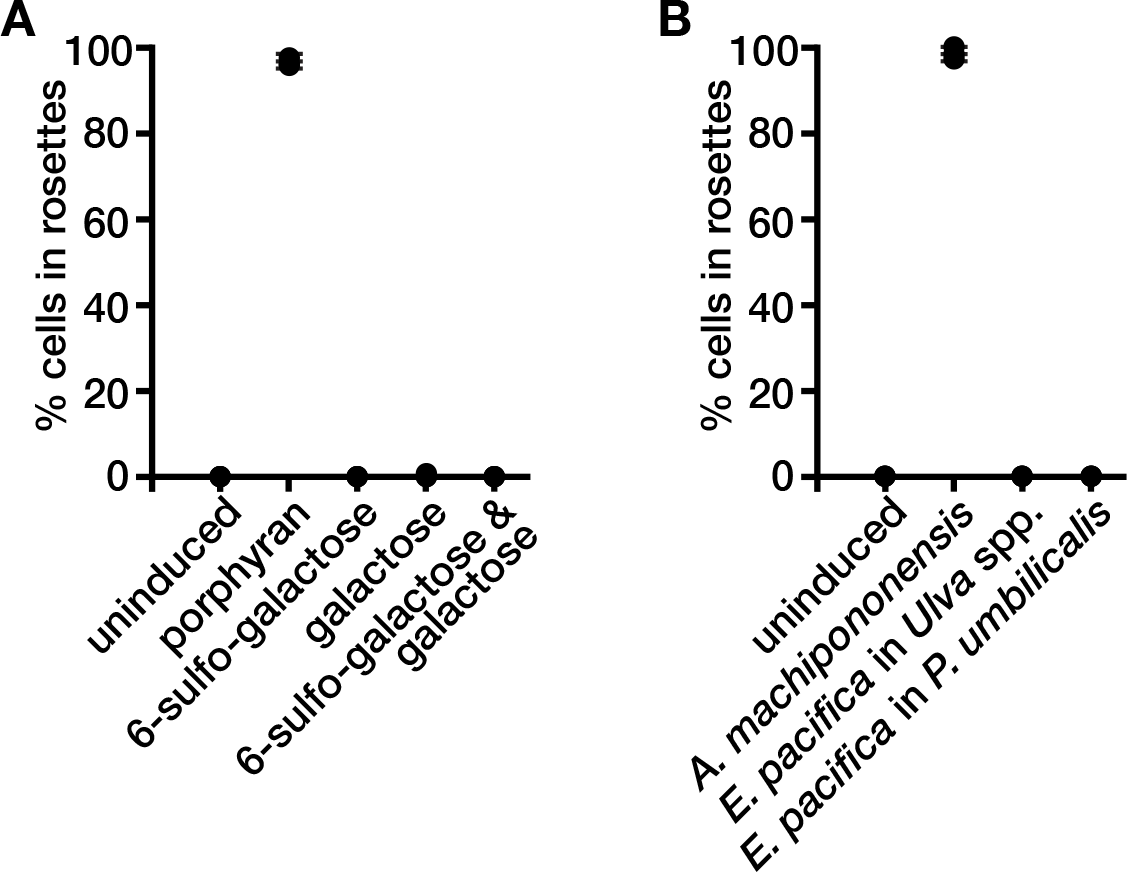
*S. rosetta* responds to porphyran, not derivative molecules. **(A)** *S. rosetta* develops into rosettes with fully intact porphyran, not its constituent monosaccharides. We added an equal mass of the indicated sugars to test if the monosaccharide D-galactose or 6-sulfo-galactose, which are alternately linked in porphyran (Figure 2C), can induce rosettes. Rosette induction was quantified as described in Figure 1B. **(B)** Outer membrane vesicles (OMVs) isolated from *E. pacifica* do not induce rosette development. We isolated OMVs from *E. pacifica* that was grown in media prepared from either *P. umbilicalis* (1% w/v steeped at 80°C, which induces rosettes) or *Ulva* spp. (not rosette inducing), and we isolated OMVs from *A. machipongonensis* grown in a media made from glycerol, yeast extract, and peptone that does not induce rosettes^2,6^. The outer membrane vesicles were resuspended to the same concentration by mass (i.e. mg/ml). Only OMVs from *A. machipongonensis* induce rosette development. This result suggests that our rosette-inducing molecule is a product of red algae, and not a secondary molecule from *E.pacifica,* our feeder bacteria.

**Figure S5:**
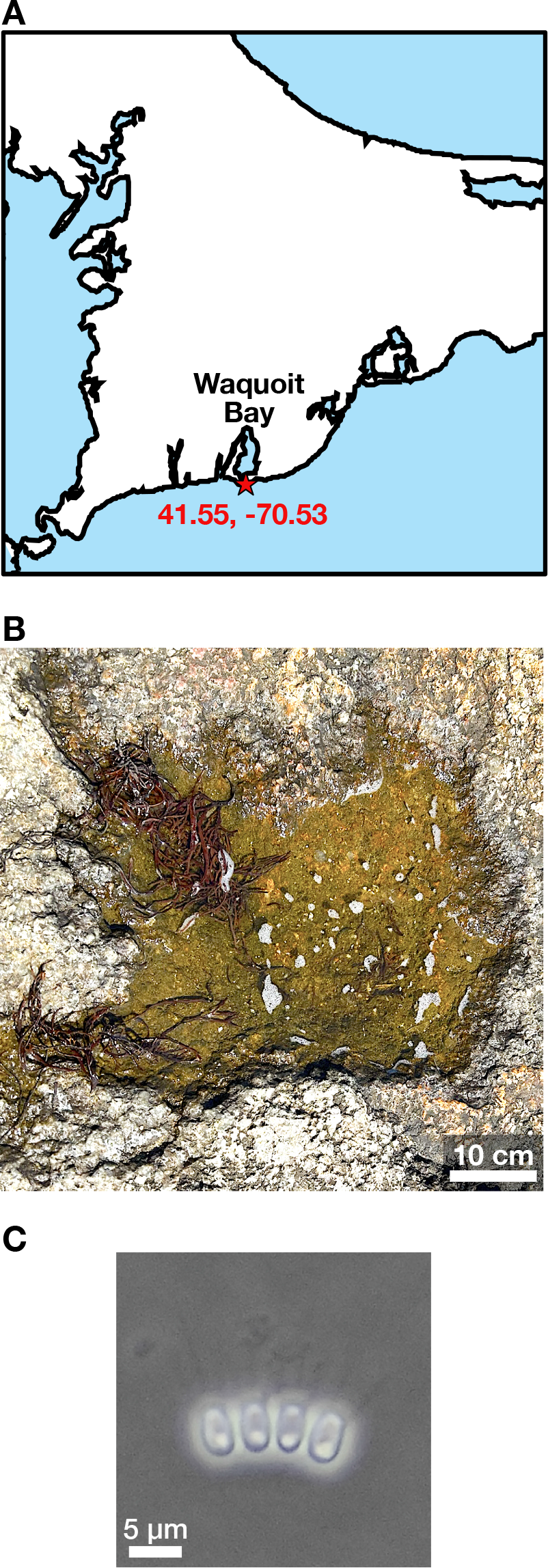
An isolated pool with fronds of red algae contains chain-forming choanoflagellates. (A) The location of a jetty at the mouth of Waquoit Bay, Massachusetts (41.55°N, 70.53°W). (B) A pool atop of a jetty at Wacquoit Bay contains fragments of red algae. The algae do not obviously anchor to the rock that forms the jetty, so waves, tides, and or storms may have swept this piece of algae on top of the jetty. (C) A chain forming choanoflagellate was abundant in the water column of the isolated pool. After collecting a sample from the jetty, we visualized it in the lab with phase contrast microscopy, and with a haemocytometer, we counted the abundance of the choanoflagellates to be ∼10^4^ cells/ml.

**Table S1:**
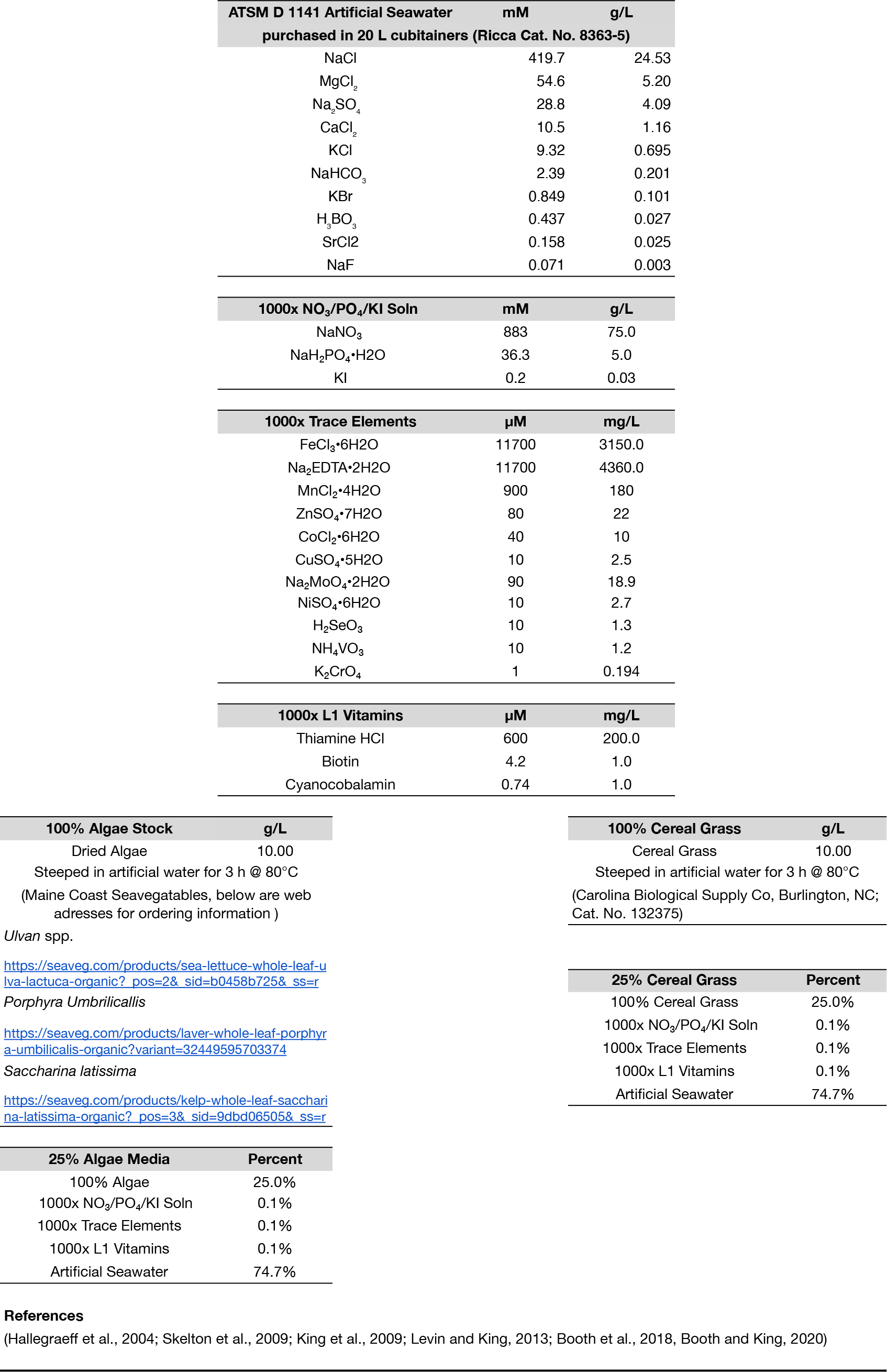
Growth Media Recipes. Media for culturing *S. rosetta* are based on seawater supplemented with nutritional enrichments. The major source of nutrition comes from cereal grass and algae steep in seawater or from peptone, yeast extract and glycerol. Additionally, solutions of vitamins, trace elements, and minerals provided critical micronutrients for *S. rosetta* growth. This table compiles all of the recipes for the stocks to prepare media.

**Table S2:**
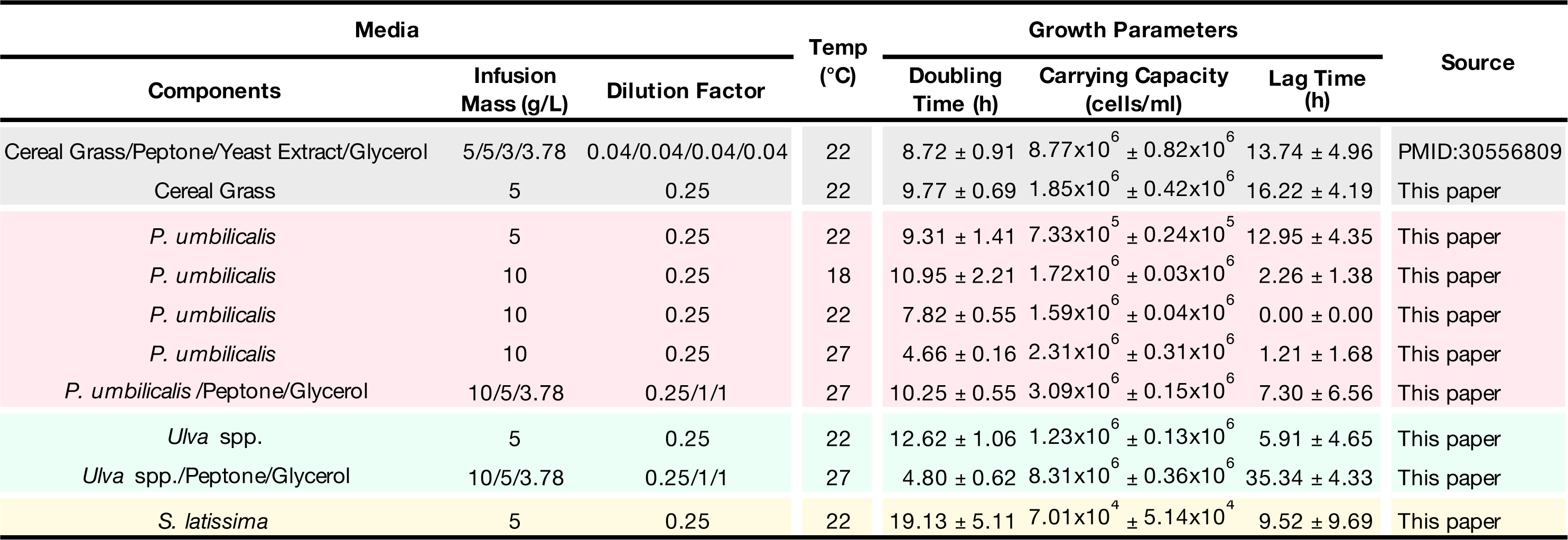
Comparison *Salpingoeca rosetta* growth parameters in algal media. The parameters calculated from time courses of *S. rosetta* growth in various media were compiled in this for ease of reference.

**Table S3:**
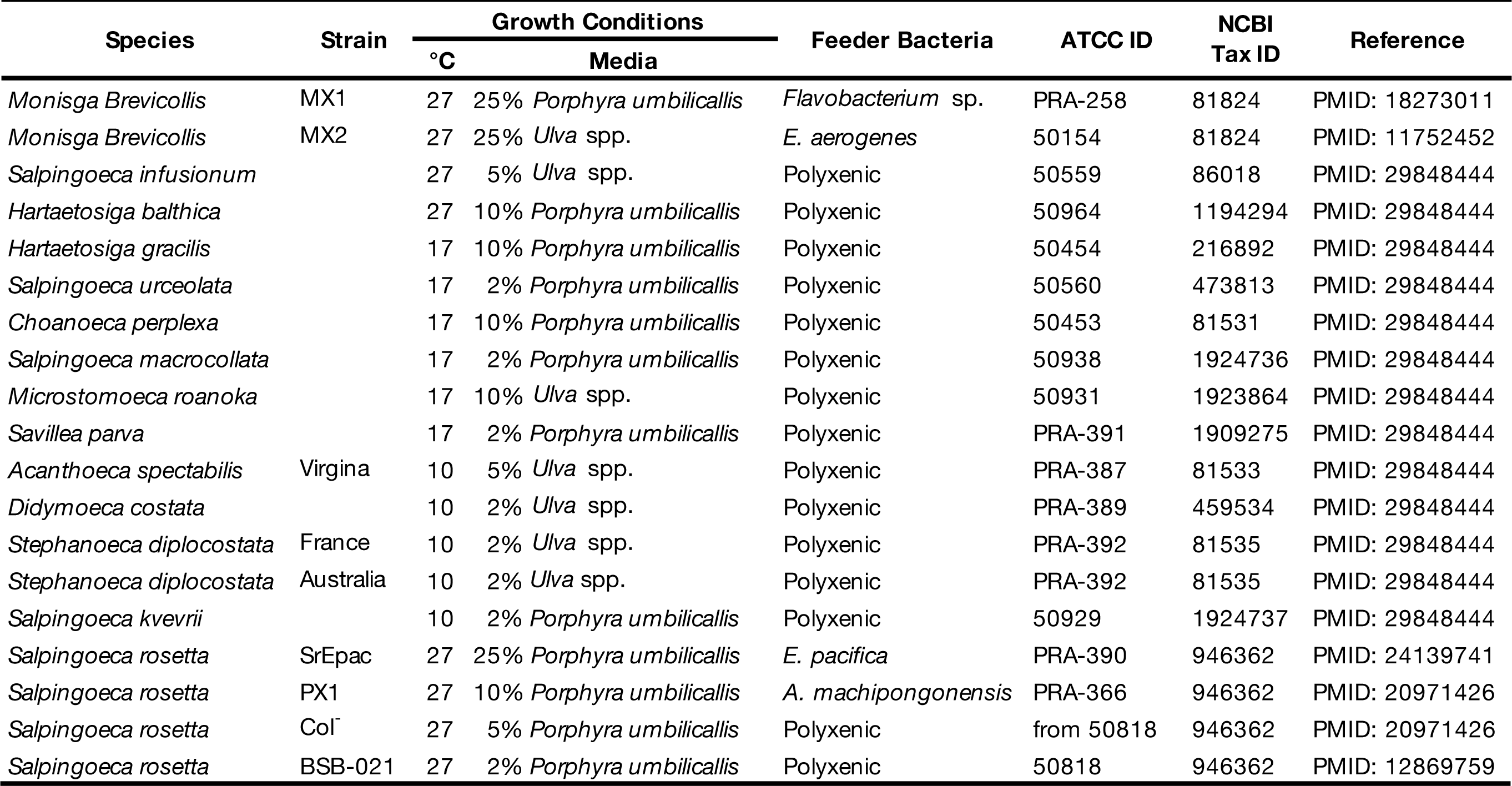
Parameters to grow diverse choanoflagellates in algae-enriched media. A variety of choanoflagellates were grown in media prepared from *P. umbilicalis* or *Ulva* spp., showing the generality of algal media to support the growth of diverse choanoflagellates.

## References

1. Dayel, M.J., Alegado, R.A., Fairclough, S.R., Levin, T.C., Nichols, S.A., McDonald, K., and King, N. (2011). Cell differentiation and morphogenesis in the colony-forming choanoflagellate Salpingoeca rosetta. Dev. Biol. 357, 73–82. 10.1016/j.ydbio.2011.06.003.

2. Levin, T.C., and King, N. (2013). Evidence for Sex and Recombination in the Choanoflagellate Salpingoeca rosetta. Curr. Biol. 23, 2176–2180. 10.1016/j.cub.2013.08.061.

3. Woznica, A., Gerdt, J.P., Hulett, R.E., Clardy, J., and King, N. (2017). Mating in the Closest Living Relatives of Animals Is Induced by a Bacterial Chondroitinase. Cell 170, 1175–1183.e11. 10.1016/j.cell.2017.08.005.

4. Ireland, E.V., Woznica, A., and King, N. (2020). Synergistic Cues from Diverse Bacteria Enhance Multicellular Development in a Choanoflagellate. Appl. Environ. Microbiol. 86, e02920–19. 10.1128/AEM.02920-19.

5. Alegado, R.A., Brown, L.W., Cao, S., Dermenjian, R.K., Zuzow, R., Fairclough, S.R., Clardy, J., and King, N. (2012). A bacterial sulfonolipid triggers multicellular development in the closest living relatives of animals. eLife 1, e00013. 10.7554/eLife.00013.

6. Woznica, A., Cantley, A.M., Beemelmanns, C., Freinkman, E., Clardy, J., and King, N. (2016). Bacterial lipids activate, synergize, and inhibit a developmental switch in choanoflagellates. Proc. Natl. Acad. Sci. U. S. A. 113, 7894–7899. 10.1073/pnas.1605015113.

7. McFall-Ngai, M., Hadfield, M.G., Bosch, T.C.G., Carey, H.V., Domazet-Lošo, T., Douglas, A.E., Dubilier, N., Eberl, G., Fukami, T., Gilbert, S.F., et al. (2013). Animals in a bacterial world, a new imperative for the life sciences. Proc. Natl. Acad. Sci. 110, 3229–3236. 10.1073/pnas.1218525110.

8. Alegado, R.A., and King, N. (2014). Bacterial Influences on Animal Origins. Cold Spring Harb. Perspect. Biol. 6, a016162. 10.1101/cshperspect.a016162.

9. Woznica, A., and King, N. (2018). Lessons from simple marine models on the bacterial regulation of eukaryotic development. Curr. Opin. Microbiol. 43, 108–116. 10.1016/j.mib.2017.12.013.

10. Woznica, A. (2024). What choanoflagellates can teach us about symbiosis. PLOS Biol. 22, e3002561. 10.1371/journal.pbio.3002561.

11. Pringsheim, E.G. (1956). Micro-Organisms from Decaying Seaweed. Nature 178, 480–481. 10.1038/178480a0.

12. Wahl, M., Goecke, F., Labes, A., Dobretsov, S., and Weinberger, F. (2012). The Second Skin: Ecological Role of Epibiotic Biofilms on Marine Organisms. Front. Microbiol. 3. 10.3389/fmicb.2012.00292.

13. Egan, S., Harder, T., Burke, C., Steinberg, P., Kjelleberg, S., and Thomas, T. (2013). The seaweed holobiont: understanding seaweed–bacteria interactions. FEMS Microbiol. Rev. 37, 462–476. 10.1111/1574-6976.12011.

14. Dang, H., and Lovell, C.R. (2015). Microbial Surface Colonization and Biofilm Development in Marine Environments. Microbiol. Mol. Biol. Rev. 80, 91–138. 10.1128/mmbr.00037-15.

15. Florez, J.Z., Camus, C., Hengst, M.B., and Buschmann, A.H. (2017). A Functional Perspective Analysis of Macroalgae and Epiphytic Bacterial Community Interaction. Front. Microbiol. 8. 10.3389/fmicb.2017.02561.

16. Lemay, M.A., Martone, P.T., Hind, K.R., Lindstrom, S.C., and Wegener Parfrey, L. (2018). Alternate life history phases of a common seaweed have distinct microbial surface communities. Mol. Ecol. 27, 3555–3568. 10.1111/mec.14815.

17. Lemay, M.A., Chen, M.Y., Mazel, F., Hind, K.R., Starko, S., Keeling, P.J., Martone, P.T., and Parfrey, L.W. (2021). Morphological complexity affects the diversity of marine microbiomes. ISME J. 15, 1372–1386. 10.1038/s41396-020-00856-z.

18. Ramírez-Puebla, S.T., Weigel, B.L., Jack, L., Schlundt, C., Pfister, C.A., and Mark Welch, J.L. (2022). Spatial organization of the kelp microbiome at micron scales. Microbiome 10, 52. 10.1186/s40168-022-01235-w.

19. Hehemann, J.-H., Correc, G., Barbeyron, T., Helbert, W., Czjzek, M., and Michel, G. (2010). Transfer of carbohydrate-active enzymes from marine bacteria to Japanese gut microbiota. Nature 464, 908–912. 10.1038/nature08937.

20. Barbeyron, T., Thomas, F., Barbe, V., Teeling, H., Schenowitz, C., Dossat, C., Goesmann, A., Leblanc, C., Oliver Glöckner, F., Czjzek, M., et al. (2016). Habitat and taxon as driving forces of carbohydrate catabolism in marine heterotrophic bacteria: example of the model algae-associated bacterium Zobellia galactanivorans DsijT. Environ. Microbiol. 18, 4610–4627. 10.1111/1462-2920.13584.

21. Lin, J.D., Lemay, M.A., and Parfrey, L.W. (2018). Diverse Bacteria Utilize Alginate Within the Microbiome of the Giant Kelp Macrocystis pyrifera. Front. Microbiol. 9. 10.3389/fmicb.2018.01914.

22. Martin, M., Barbeyron, T., Martin, R., Portetelle, D., Michel, G., and Vandenbol, M. (2015). The Cultivable Surface Microbiota of the Brown Alga Ascophyllum nodosum is Enriched in Macroalgal-Polysaccharide-Degrading Bacteria. Front. Microbiol. 6. 10.3389/fmicb.2015.01487.

23. Miranda, L.N., Hutchison, K., Grossman, A.R., and Brawley, S.H. (2013). Diversity and Abundance of the Bacterial Community of the Red Macroalga Porphyra umbilicalis: Did Bacterial Farmers Produce Macroalgae? PLOS ONE 8, e58269. 10.1371/journal.pone.0058269.

24. Armstrong, E., Rogerson, A., and Leftley, J.W. (2000). The Abundance of Heterotrophic Protists Associated with Intertidal Seaweeds. Estuar. Coast. Shelf Sci. 50, 415–424. 10.1006/ecss.1999.0577.

25. 25. Brocks, J.J., Nettersheim, B.J., Adam, P., Schaeffer, P., Jarrett, A.J.M., Güneli, N., Liyanage, T., van Maldegem, L.M., Hallmann, C., and Hope, J.M. (2023). Lost world of complex life and the late rise of the eukaryotic crown. Nature 618, 767–773. 10.1038/s41586-023-06170-w.

26. Brocks, J.J., Jarrett, A.J.M., Sirantoine, E., Hallmann, C., Hoshino, Y., and Liyanage, T. (2017). The rise of algae in Cryogenian oceans and the emergence of animals. Nature 548, 578–581. 10.1038/nature23457.

27. 27. Manual of the infusoria, including a description of all known flagellate, ciliate, and tentaculiferous protozoa, British and foreign and an account of the organization and affinities of the sponges by W. Saville Kent, 3 vol (1881).

28. Leadbeater, B.S.C., and Thomsen, H.A. (2000). Choanoflagellida. In The Illustrated Guide to the Protozoa (Allen Press Inc.), pp. 14–38.

29. 29. Leadbeater, B.S.C. (2015). The choanoflagellates: evolution, biology, and ecology (Cambridge University Press).

30. Booth, D.S., and King, N. (2020). Genome editing enables reverse genetics of multicellular development in the choanoflagellate Salpingoeca rosetta. eLife 9, e56193. 10.7554/eLife.56193.

31. Croft, M.T., Lawrence, A.D., Raux-Deery, E., Warren, M.J., and Smith, A.G. (2005). Algae acquire vitamin B12 through a symbiotic relationship with bacteria. Nature 438, 90–93. 10.1038/nature04056.

32. Wells, M.L., Potin, P., Craigie, J.S., Raven, J.A., Merchant, S.S., Helliwell, K.E., Smith, A.G., Camire, M.E., and Brawley, S.H. (2017). Algae as nutritional and functional food sources: revisiting our understanding. J. Appl. Phycol. 29, 949–982. 10.1007/s10811-016-0974-5.

33. Brawley, S.H., Blouin, N.A., Ficko-Blean, E., Wheeler, G.L., Lohr, M., Goodson, H.V., Jenkins, J.W., Blaby-Haas, C.E., Helliwell, K.E., Chan, C.X., et al. (2017). Insights into the red algae and eukaryotic evolution from the genome of Porphyra umbilicalis (Bangiophyceae, Rhodophyta). Proc. Natl. Acad. Sci. 114, E6361–E6370. 10.1073/pnas.1703088114.

34. Lee, J.J., and Soldo, A.T. (Anthony T.) (1992). Protocols in protozoology (Society of Protozoologists).

35. King, N., Young, S.L., Abedin, M., Carr, M., and Leadbeater, B.S.C. (2009). Starting and Maintaining Monosiga brevicollis Cultures. Cold Spring Harb. Protoc. 2009, pdb.prot5148. 10.1101/pdb.prot5148.

36. Levin, T.C., Greaney, A.J., Wetzel, L., and King, N. (2014). The Rosetteless gene controls development in the choanoflagellate S. rosetta. eLife 3. 10.7554/eLife.04070.

37. Booth, D.S., and King, N. (2020). Genome editing enables reverse genetics of multicellular development in the choanoflagellate Salpingoeca rosetta. eLife 9, e56193. 10.7554/eLife.56193.

38. Fairclough, S.R., Dayel, M.J., and King, N. (2010). Multicellular development in a choanoflagellate. Curr. Biol. 20, R875–R876. 10.1016/j.cub.2010.09.014.

39. Peng, C.-C., Dormanns, N., Regestein, L., and Beemelmanns, C. (2023). Isolation of sulfonosphingolipids from the rosette-inducing bacterium Zobellia uliginosa and evaluation of their rosette-inducing activity. RSC Adv. 13, 27520–27524. 10.1039/D3RA04314B.

40. Mukherjee, S., Seshadri, R., Varghese, N.J., Eloe-Fadrosh, E.A., Meier-Kolthoff, J.P., Göker, M., Coates, R.C., Hadjithomas, M., Pavlopoulos, G.A., Paez-Espino, D., et al. (2017). 1,003 reference genomes of bacterial and archaeal isolates expand coverage of the tree of life. Nat. Biotechnol. 35, 676–683. 10.1038/nbt.3886.

41. Droop, M.R., and Doyle, J. (1966). Ubiquinone as a Protozoan Growth Factor. Nature 212, 1474–1475. 10.1038/2121474a0.

42. Brunet, T., Larson, B.T., Linden, T.A., Vermeij, M.J.A., McDonald, K., and King, N. (2019). Light-regulated collective contractility in a multicellular choanoflagellate. Science 366, 326–334. 10.1126/science.aay2346.

43. 43. Ros-Rocher, N., Reyes-Rivera, J., Foroughijabbari, Y., Combredet, C., Larson, B.T., Coyle, M.C., Houtepen, E.A.T., Vermeij, M.J.A., King, N., and Brunet, T. (2024). Mixed clonal-aggregative multicellularity entrained by extreme salinity fluctuations in a close relative of animals. Preprint at bioRxiv, 10.1101/2024.03.25.586565.

44. Shibl, A.A., Isaac, A., Ochsenkühn, M.A., Cárdenas, A., Fei, C., Behringer, G., Arnoux, M., Drou, N., Santos, M.P., Gunsalus, K.C., et al. (2020). Diatom modulation of select bacteria through use of two unique secondary metabolites. Proc. Natl. Acad. Sci. 117, 27445–27455. 10.1073/pnas.2012088117.

45. Shemi, A., Alcolombri, U., Schatz, D., Farstey, V., Vincent, F., Rotkopf, R., Ben-Dor, S., Frada, M.J., Tawfik, D.S., and Vardi, A. (2021). Dimethyl sulfide mediates microbial predator–prey interactions between zooplankton and algae in the ocean. Nat. Microbiol. 6, 1357–1366. 10.1038/s41564-021-00971-3.

46. Becker, S., Tebben, J., Coffinet, S., Wiltshire, K., Iversen, M.H., Harder, T., Hinrichs, K.-U., and Hehemann, J.-H. (2020). Laminarin is a major molecule in the marine carbon cycle. Proc. Natl. Acad. Sci. 117, 6599–6607. 10.1073/pnas.1917001117.

47. Vidal-Melgosa, S., Sichert, A., Francis, T.B., Bartosik, D., Niggemann, J., Wichels, A., Willats, W.G.T., Fuchs, B.M., Teeling, H., Becher, D., et al. (2021). Diatom fucan polysaccharide precipitates carbon during algal blooms. Nat. Commun. 12, 1150. 10.1038/s41467-021-21009-6.

48. Sanz, V., Torres, M.D., Domínguez, H., Pinto, I.S., Costa, I., and Guedes, A.C. (2023). Seasonal and spatial compositional variation of the red algae Mastocarpus stellatus from the Northern coast of Portugal. J. Appl. Phycol. 35, 419–431. 10.1007/s10811-022-02863-3.

49. Lipinska, A.P., Collén, J., Krueger-Hadfield, S.A., Mora, T., and Ficko-Blean, E. (2020). To gel or not to gel: differential expression of carrageenan-related genes between the gametophyte and tetasporophyte life cycle stages of the red alga Chondrus crispus. Sci. Rep. 10, 11498. 10.1038/s41598-020-67728-6.

50. Gf, B., Jm, G., P, R., Ph, G., and F, M. (1996). Quantifying the short-term temperature effect on light-saturated photosynthesis of intertidal microphytobenthos. Mar. Ecol. Prog. Ser. 134, 309–313. 10.3354/meps134309.

51. Noll, P., Lilge, L., Hausmann, R., and Henkel, M. (2020). Modeling and Exploiting Microbial Temperature Response. Processes 8, 121. 10.3390/pr8010121.

52. Dowd, W.W., King, F.A., and Denny, M.W. (2015). Thermal variation, thermal extremes and the physiological performance of individuals. J. Exp. Biol. 218, 1956–1967. 10.1242/jeb.114926.

53. Torres, P.B., Nagai, A., Jara, C.E.P., Santos, J.P., Chow, F., and Santos, D.Y.A.C. dos (2021). Determination of sulfate in algal polysaccharide samples: a step-by-step protocol using microplate reader. Ocean Coast. Res. 69, e21021.

54. Hallegraeff, G.M., Anderson, D.M., Cembella, A.D., and Enevoldsen, H.O. (2004). Manual on harmful marine microalgae (UNESCO).

55. 55. Leon, F., Esparza, J., Deng, V., Coyle, M.C., and Espinoza, S. (2024). Cell-type-specific gene expression enables the choanoflagellate S. rosetta to utilize of colloidal iron. unpublished.

56. Symons, M.H., and Mitchison, T.J. (1991). Control of actin polymerization in live and permeabilized fibroblasts. J. Cell Biol. 114, 503–513. 10.1083/jcb.114.3.503.

57. Cramer, L.P., and Mitchison, T.J. (1995). Myosin is involved in postmitotic cell spreading. J. Cell Biol. 131, 179–189. 10.1083/jcb.131.1.179.

58. 58. Merkle, R.K., and Poppe, I. (1994). [1] Carbohydrate composition analysis of glycoconjugates by gas-liquid chromatography/mass spectrometry. In Methods in Enzymology Guide to Techniques in Glycobiology. (Academic Press), pp. 1–15. 10.1016/0076-6879(94)30003-8.

59. Heiss, C., Wang, Z., Thurlow, C.M., Hossain, M.J., Sun, D., Liles, M.R., Saper, M.A., and Azadi, P. (2019). Structure of the capsule and lipopolysaccharide O-antigen from the channel catfish pathogen, Aeromonas hydrophila. Carbohydr. Res. 486, 107858. 10.1016/j.carres.2019.107858.

